# Population-level survey of loss-of-function mutations revealed that background dependent fitness genes are rare and functionally related in yeast

**DOI:** 10.1101/2021.08.25.457624

**Authors:** Elodie Caudal, Anne Friedrich, Arthur Jallet, Marion Garin, Jing Hou, Joseph Schacherer

## Abstract

In natural populations, the same mutation can lead to different phenotypic outcomes due to the genetic variation that exists among individuals. Such genetic background effects are commonly observed, including in the context of many human diseases. However, systematic characterization of these effects at the species level is still lacking to date. Here, we sought to comprehensively survey background-dependent traits associated with gene loss-of-function (LoF) mutations in 39 natural isolates of *Saccharomyces cerevisiae* using a transposon saturation strategy. By analyzing the modeled fitness variability of a total of 4,469 genes, we found that 15% of them, when impacted by a LoF mutation, exhibited a significant gain- or loss-of-fitness phenotype in certain natural isolates compared to the reference strain S288C. Out of these 632 genetic background-dependent fitness genes identified, a total of 2/3 show a continuous variation across the population while 1/3 are specific to a single genetic background. Genes related to mitochondrial function are significantly overrepresented in the set of genes showing a continuous variation and display a potential functional rewiring with other genes involved in transcription and chromatin remodeling as well as in nuclear-cytoplasmic transport. Such rewiring effects are likely modulated by both the genetic background and the environment. While background-specific cases are rare and span diverse cellular processes, they can be functionally related at the individual level. All background-dependent fitness genes tend to have an intermediate connectivity in the global genetic interaction network and have shown relaxed selection pressure at the population level, highlighting their potential evolutionary characteristics.

## Introduction

The same mutation might show different phenotypic effects across genetically distinct individuals due to standing genomic variation^1–8^. Such background effects have been described across species and impact the phenotype-genotype relationship, including in the context of health and disease. Indeed, they have been observed in multiple human Mendelian disorders, where individuals carrying the same causal mutation can display a wide range of clinical symptoms, including variable severity and age-of-onset^1,7,9–12^. The underlying origin of these background effects may be both intrinsic, *i*.*e*. due to interactions between the causal variant and other genetic modifiers^9–11^ and/or extrinsic, *i*.*e*. due to environmental factors^12,13^. To date, a handful of modifier genes have been found associated with human disorders, most notably in cystic fibrosis^11,12^. However, such examples remain rare and anecdotal due to the low number of sample cases in most human Mendelian diseases.

In recent years, several large-scale surveys in different model organisms such as the yeasts *Saccharomyces cerevisiae* and *Schizosaccharomyces pombe*, the nematode *Caenorhabditis elegans* and various human cell lines highlighted the broad influence of genetic backgrounds on the phenotypic outcomes associated with loss-of-function mutations^14–22^. In yeast, a study comparing systematic gene deletion collections in two laboratory strains, Σ1278b and S288C, showed that approximately 1% of all genes (57/5100) can display background-dependent gene essentiality, *i*.*e*. where the deletion of the same gene can be lethal in one background but not the other^14^. Several origins underlying such gene essentiality have been identified, including genetic interactions between the mitochondrial genome and/or viral elements with the nuclear genome^23^ as well as genetic interactions between the primary deletion gene and background-specific modifiers^24^. While gene essentiality may be the most severe manifestation associated with loss-of-function mutations, gain- and loss-of-fitness variation related to genetic backgrounds or environmental conditions were also found in yeast^5,25^. For example, about 20% of yeast genes showed background-dependent fitness variation under a wide range of growth conditions, including the presence of various drugs, osmotic stress, and nutrient sources in 4 genetically diverse isolates^25^. However, all these studies only include a limited number of genetic backgrounds and therefore cannot accurately reflect the extend of the background effect at the species level.

Recently, a large collection of 1,011 *S. cerevisiae* isolates originated from various ecological and geographical sources has been completely sequenced^26^, representing an incomparable resource to systematically study the effects of genetic backgrounds at the species level. Several strategies have been developed in *S. cerevisiae* to explore the impact of loss-of-function mutations, including systematic gene deletions using homologous recombination^14^, gene-disruption using the CRISPR-Cas9 editing systems^27^, repeated backcrosses^25^ and transposon mutagenesis^28–30^. Among these strategies, transposon mutagenesis based on random excision and insertions are particularly attractive for exploring in parallel a large number of genetically diverse individuals. This method relies on transposition events via a carrier plasmid, which allow for the generation of millions of mutants carrying genomic insertions leading to loss-of-function mutations^30^. Due to the random insertion patterns in each genetic background, these methods do not depend on sequence homology as is the case for traditional PCR-based gene deletions and CRISPR-Cas9 related strategies^27,31^, and they do not present the risk of inadvertently introducing exogenous genomic regions as it might be the case for backcross-based strategies^25^.

Here, we selected over a hundred natural isolates broadly representative of the diversity *S. cerevisiae* species, and performed transposon saturation analyses using the *Hermes* transposition system^29^. We generated, sequenced and analyzed large pools of transposon insertion mutants and constructed a logistic model to predict the fitness effects of gene loss-of-function based on the insertion densities in and around of each annotated gene. Comparing the fitness prediction between the different isolates and the S288C reference, we identified 632 background-dependent fitness genes, corresponding to approximately 15% of the genome. Overall, they are functionally related, with members of the same protein complex of biological process showing similar variability in each genetic background. They also tend to show an intermediate level of integration in genetic networks compared to non-essential and essential genes, and might be under positive or relaxed purifying selection at the population level.

## Results

### Generation of LoF mutant collections using the *Hermes* transposon system

To gain insight into fitness variation associated with loss-of-function mutations across different *S. cerevisiae* genetic backgrounds, we performed transposon saturation assays in various natural isolates using the *Hermes* transposon system. The *Hermes* transposon system has previously been adapted in yeast to allow the selection of random insertion events in liquid culture, which makes this system particularly suitable for parallel analyses of a large number of genetically diverse individuals^29^. This system is based on a centromeric plasmid, which contains the *Hermes* transposase under the control of a modified galactose inducible promoter *GalS*, as well as a transposon carrying a selectable marker (Figure 1A). Briefly, for any strain of interest, the plasmid is first transformed into stable haploid cells and then propagated in media containing galactose to induce excision and reinsertion of the transposon at random locations in the genome, thereby generating a large pool of individuals with hundreds of thousands of insertions along the genome (Figure 1A). After a recovery phase in rich medium, the genome of this pool of mutants is recovered, then fragmentated and circularized (Figure 1A). Using PCR with outward facing primers specifically targeting the transposon, a library that exclusively contains the insertion sites can be constructed and then sequenced using standard Illumina methods (Figure 1A). In principle, transposon insertions that cause severe fitness defects, for example those occurring in essential genes, will not be recovered due to the competitive disadvantage compared to events occurring in genes which are not essential. Analysis of insertion patterns along the genomes of different individuals therefore provides a proxy for fitness variation related to loss-of-function mutations.

**Figure 1.**
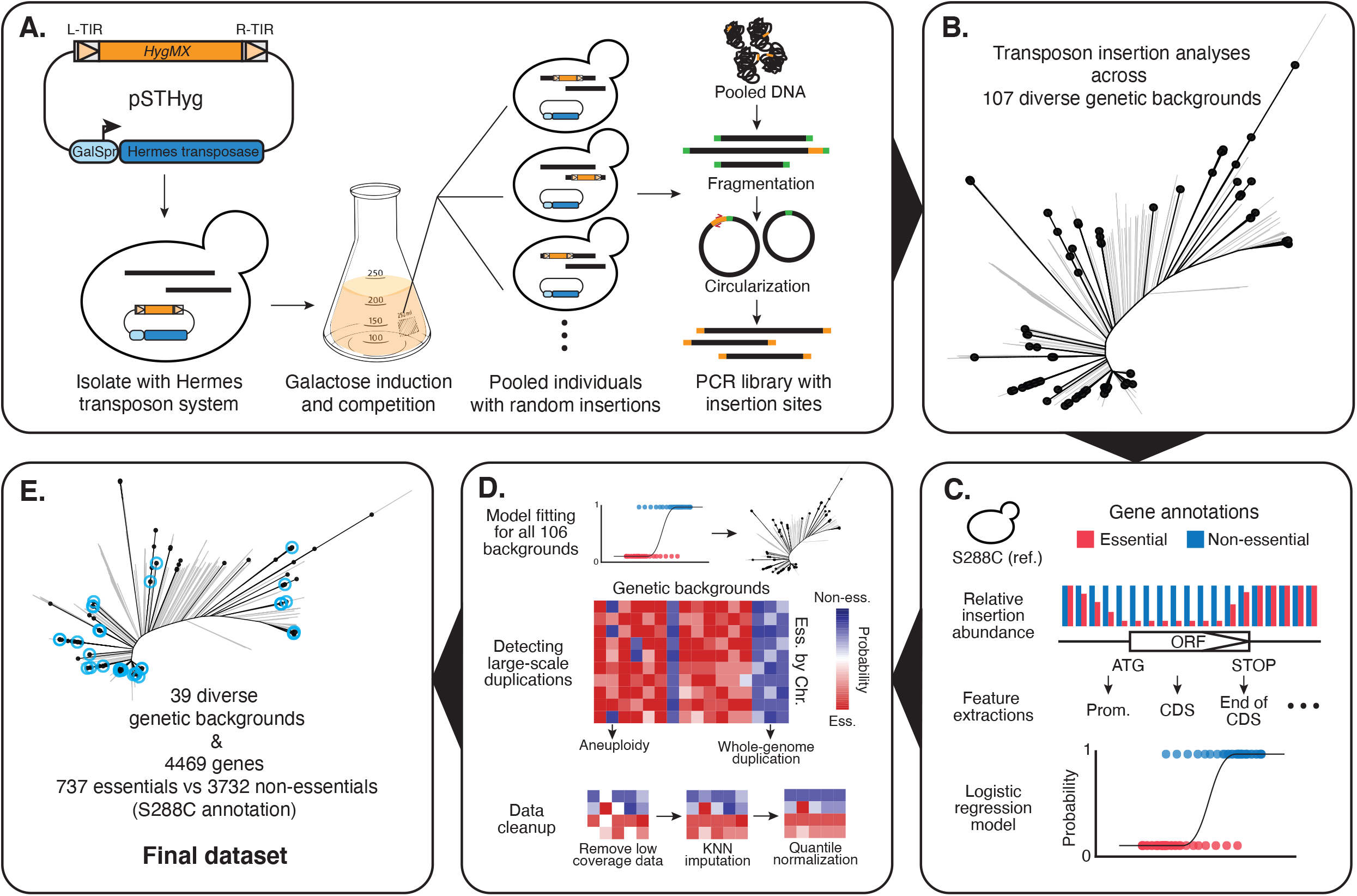
Summary of the *Hermes* transposon saturation procedure. (A) A centromeric plasmid carrying the *Hermes* transposase and a transposon containing a hygromycin resistance marker (*HygMX*) is transformed into a haploid isolate background. Random transposon insertions are induced and selected. The mutant pool is then recovered and a PCR library that contains only the insertion sites is constructed and sequence. (B) Distribution of the selected 107 isolates across the species. The neighbour-joining tree was constructed using biallelic SNPs in the 1,011 yeast collection^26^. Selected strains are highlighted in black. (C) A logistic model was constructed using insertion profiles in the reference strain S288C. Gene essentiality annotations were used as a binary classifier, excluding those annotated as involved in galactose metabolism, respiration and slow growth. (D) The logistic model was applied to insertion patterns in the remaining 106 isolates. Large-scale genome duplications were detected by looking at fitness predictions for all annotated essential genes along each chromosome. Low coverage regions were removed then imputed using k-nearest-neighbour method. The imputed fitness matrix was then quantile normalized. (E) The final dataset after imputation consists of 39 isolates and 4469 genes. Strains included in the final dataset are highlighted in blue.

In addition to the S288C reference strain, we selected 106 isolates originated from various ecological and geographical sources that are broadly representative of the species diversity (Figure 1B, Table S1). Stable haploid variants of this set of isolates have been generated previously^32,33^ and are all capable of growing in galactose medium. We have adapted the initial version of the *Hermes* transposon plasmid to carry a hygromycin resistance marker instead of nourseothricin to ensure compatibility with selected strains, which may carry either a *KanMX* or *NatMX* marker at the *HO* locus. Transposon insertion profiles for each isolate were obtained as described (Figure 1A). We observed a marked variability in terms of insertion efficiency across different genetic backgrounds, ranging from ∼100 to ∼300,000 unique insertion sites (Figure S1A, Table S1). No discernible correlation between the genetic origin of the isolates and the transposon insertion efficiency was observed (Table S1). We then compared the insertion preferences between the S288C reference strain and the 106 natural isolates (Figure S1B). Insertion densities for known sequence motifs^29^ were conserved across the different genetic backgrounds (Figure S1B).

Using insertion profiles and the annotation of gene essentiality in the S288C reference, we analyzed the average insertion patterns in the promoters (−500 bp to ATG, 100 bp window), the coding DNA sequences (CDS), and the terminators (STOP to +500 bp, 100 bp window) for all annotated essential *vs*. non-essential genes and found that the number of insertion drops from -100 bp prior to CDS and extends to - 100 bp until the terminator region, with on average ∼3 times fewer insertions within the CDS in the essential genes compared to non-essential genes (Figure S1C-E). This pattern is consistent with the results obtained in previous studies using the *Hermes* system^29^.

### Modeling based on insertion patterns to identify background-dependent fitness genes

We constructed a logistic model that simultaneously takes into account transposon insertions that have occurred in the genes and surrounding regions using the insertion profiles from the reference strain S288C and the corresponding gene essentiality annotations as a binary classifier (Figure 1C, Table S2, See Methods). We applied this model to the insertion profiles of all 107 diverse isolates (Figure 1D). For each annotated ORF (approximately a total of 6,300 ORFs), a probability was calculated based on the model, ranging from a value of 1, corresponding to most likely non-essential, to 0, corresponding to most likely essential. Genomic regions with low insertion densities contribute to overall low predictive powers (Figure S2A-B), which were subsequently removed. Due to the variability of insertion efficiency across strains (Figure S1A), removal of regions with low insertion density has led to entire strain backgrounds with few interpretable genes. By maximizing both the number of strains and genes that remained after data imputation (Figure S2C-D), a total of 52 backgrounds and probability predictions for 4,469 genes were retained for subsequent analyses (Table S3).

Large-scale genome duplications, including aneuploidies and endoreduplications, are frequently observed in yeast experimental evolution^34–36^. Such events may hamper the accuracy of the modeled fitness effect in the context of the transposon insertion assay, as genes in the duplicated region will all appear to be fit due to insertions in a single copy of the gene. We searched for signals of large-scale genome duplications by examining all annotated essential genes along the chromosomes in our set of isolates (Figure 1D, Figure S3A). We detected endoreduplication events in 7 out of 52 strains where all chromosomes appeared to be duplicated based on the high predicted probability values for all essential genes (Figure S3A). In another set of 6 strains, essential genes showed an intermediate to high probability prediction but not high enough to be confidently classified as non-essential. These 6 strains were then confirmed as a mixture of haploid and diploid cells using flow cytometry. In addition to these whole genome events, we also detected 3 strains with an aneuploidy of chromosome I (ACT, BKL and ACV), one strain with an aneuploidy of chromosome XII (CPG) and one strain with an aneuploidy of chromosome XIV (CQA). These aneuploidies were not present in the original isolate, with the exception of chromosome I aneuploidies in ACT and ACV strains, highlighting the dynamics of genome instability in different genetic backgrounds. All 13 strains with whole genome endoreduplication were entirely removed from the dataset. We also excluded aneuploid chromosomes from the analysis (Table S3).

Next, we looked specifically at the probability predictions in the reference S288C. The final set of 4,469 genes includes 3,732 and 737 that were annotated non-essential and essential, respectively. Among the genes annotated as non-essential, approximately 180 were predicted to be likely essential in our data (Table S3), of which more than 70% correspond to slow grow or galactose-specific fitness defect genes. For example, the hexose transporters *HXT6/7* and genes involved in galactose metabolism are all predicted to be likely essential, as expected by using our transposition saturation strategy (Figure S3B). On the other hand, 26 genes annotated as essential were predicted to be likely non-essential, with a predicted probability > 0.8 (Table S3). Among these, we found *FUR1, HIP1* and *SSY5*, consisting of amino acid transporters that are only essential in the multi-auxotrophic BY4741 background, isogenic to the prototrophic S288C we used in our study (Table S3). We have also found genes where the essentiality concerns only part of the ORF, *i*.*e*. the essential domains, as has also been observed in previous studies using this transposon saturation strategy^28^ (Figure S3C). Notably, we found the *RET2* and *SRP14* genes, which are also among the essential genes specific to the S288C background compared to Σ1278b in systematic gene deletion collections^14^. Indeed, these domain essential effects are recaptured in our dataset when comparing insertion patterns between S288C and Σ1278b (Figure S3D). In fact, background-specific essential genes between S288C and Σ1278b that did not display severe fitness defect when deleted in the non-essential background^24^, including S288C-specific essential genes (*RET2, UBC1* and *SRP14*), and Σ1278b-specific essential genes (*SKI8, TMA108* and *AAT2*), all showed domain essential effects and are all recaptured in our data (Figure S3D).

Overall, the predicted probability based on our logistic model can serve as a reasonable proxy for fitness variation related to loss-of-function mutations. Modeled fitness (predicted probability for non-essentiality) is more accurate at predicting non-essential/high fitness cases than essential/low fitness cases, which may in part due to that non-essential genes are *de facto* slow growers in the context of our experimental conditions, and in part due to bias in transposon insertion densities across genomic regions and genetic backgrounds. Essentialities related to specific domains can be recaptured by the raw insertion patterns but not by our modeled fitness values (Figure S3C-D). However, this effect is inherent to the transposon saturation system and should not lead to differential fitness effect prediction in different genetic backgrounds. Our final dataset consists of 39 isolates from various origins and predicted fitness for 4,469 genes, which is analysed in more detail (Figure 1E, Table S3).

### Environmental dependency of fitness variability associated with LoF mutations across backgrounds

We first performed a hierarchical clustering based on the predicted fitness values of 4,469 genes across the 39 genetic backgrounds (Figure 2). Profile similarity based on the predicted fitness effects did not correlate with the genetic origins of the isolates (Figure 2). Genes that are consistently essential in different isolates clustered together and are enriched for essential biological processes, including ribosome biogenesis, rRNA processing, DNA replication, protein transport and cell cycle (Figure 2). Genes that are consistently non-essential in all backgrounds formed a large cluster without significant enrichment for any specific biological process. Interestingly, several clusters of genes with variable fitness effects were identified, displaying modular switches from fit to non-fit phenotypes across the entire population. Gene enrichment analyses revealed genes involved in mitochondrial translation, transcription regulation and general translational processes (Figure 2). A large proportion of these genes with population-wide fitness variation consists of nuclear encoded mitochondrial genes involved in respiration, which were expected to show a selective disadvantage in our pool of mutants that must grow on galactose. This observation suggests that such general fitness variability may be environment-related rather than background-specific *per se*. However, other biological processes in addition to respiration and mitochondrial functions have also been enriched, for which the impact of environment *vs*. genetic background on their fitness variability remains unclear.

**Figure 2.**
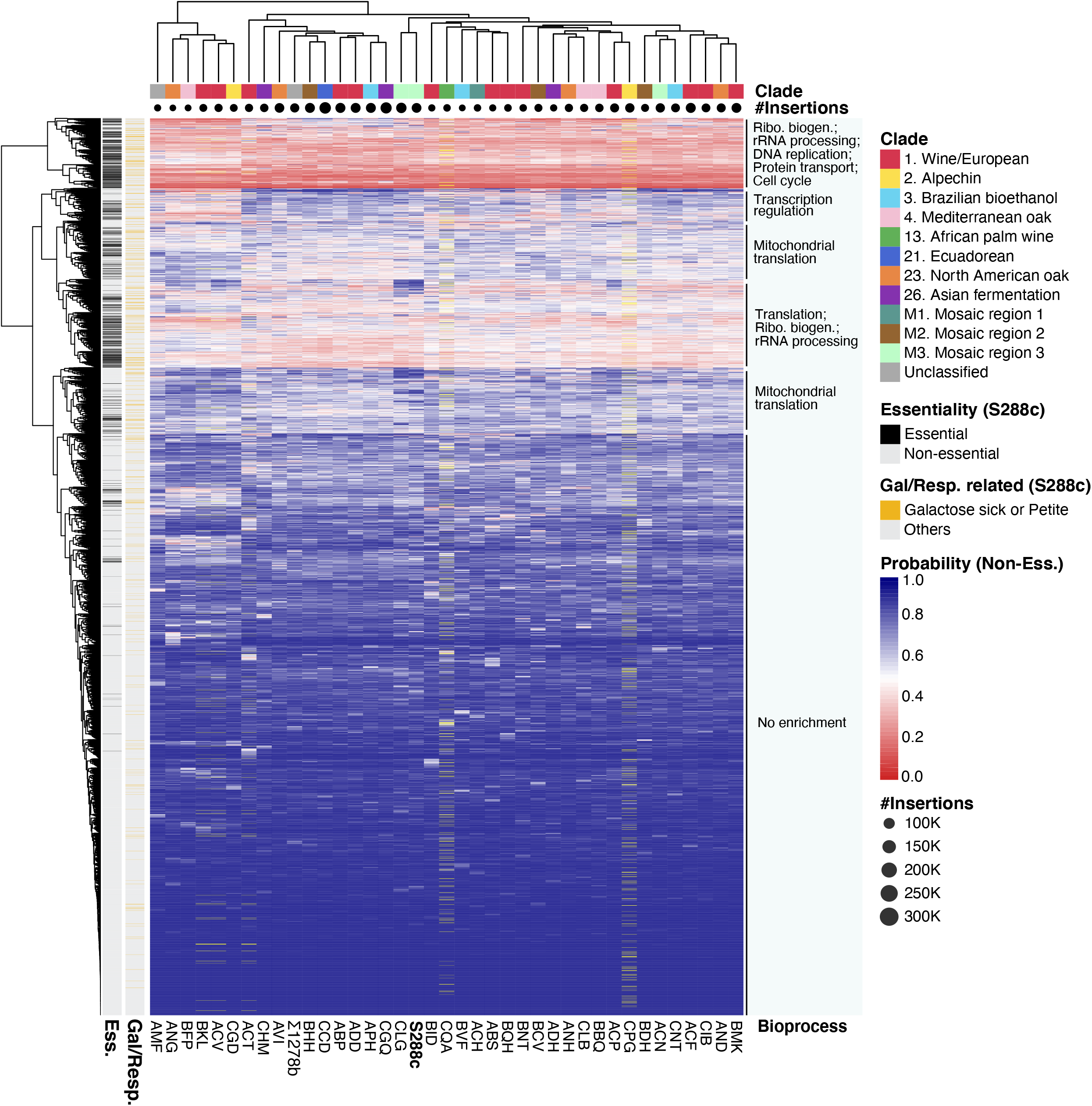
Hierarchical clustering of 4469 fitness predictions across 39 genetic backgrounds. The distance matrix was calculated using the Euclidean distance method. The genetic origin of each isolate was color-coded, and the total insertion numbers per isolate was represented by dot size under the origin color code. Genes annotated as essential in the reference S288C are highlighted in black, and genes annotated as either galactose or respiration related are highlighted in yellow on the sidebars. Genes within duplicated chromosomes were removed (yellow bars on heatmap). Biological processes that are enrichment in subclusters are annotated.

To further characterize the background-dependent fitness variation, we systematically compared the predicted fitness values for each gene in a given isolate with the predictions of the reference strain S288C. A differential fitness score for each gene in each background was calculated by subtracting the predicted fitness value in a given strain from the corresponding fitness prediction in the reference S288C. A minimum absolute value of the differential fitness score of 0.5 was considered significant, which corresponds to a *bona fide* reverse in the direction of being predicted as essential or non-essential according to our logistic model. In total, 632 genes were identified with marked fitness variation, with 458 and 174 showing a loss-of-fitness (S288C healthy and background sick) and a gain-of-fitness (S288C sick and background healthy) compared to the reference, respectively. The number of identified differential fitness genes ranges from 8 (ACP) to 88 (BQH) for loss-of-fitness cases (with a median of 61), and from 6 (CGD) to 42 (AMF) for gain-of-fitness cases (with a median of 16) (Figure 3A). A total of 163 out of all 632 hits are related to respiration and mitochondrial functions, representing ∼20% to ∼60% of loss-of-fitness hits depending on the genetic background (Figure 3A). Furthermore, these respiration-related genes tend to impact more backgrounds on average than non-respiration related hits (Figure 3B). These observations echoed what was shown on hierarchical clustering where mitochondrial related genes were highly enriched in clusters with modular fitness variation in several backgrounds (Figure 2). Again, due to the overrepresentation of these respiration-related genes and their continuous fitness variation in the population, we suspect that these hits are likely to be impacted by the environment (*i*.*e*. pooled competition in galactose media) in addition to any specific genetic backgrounds.

**Figure 3.**
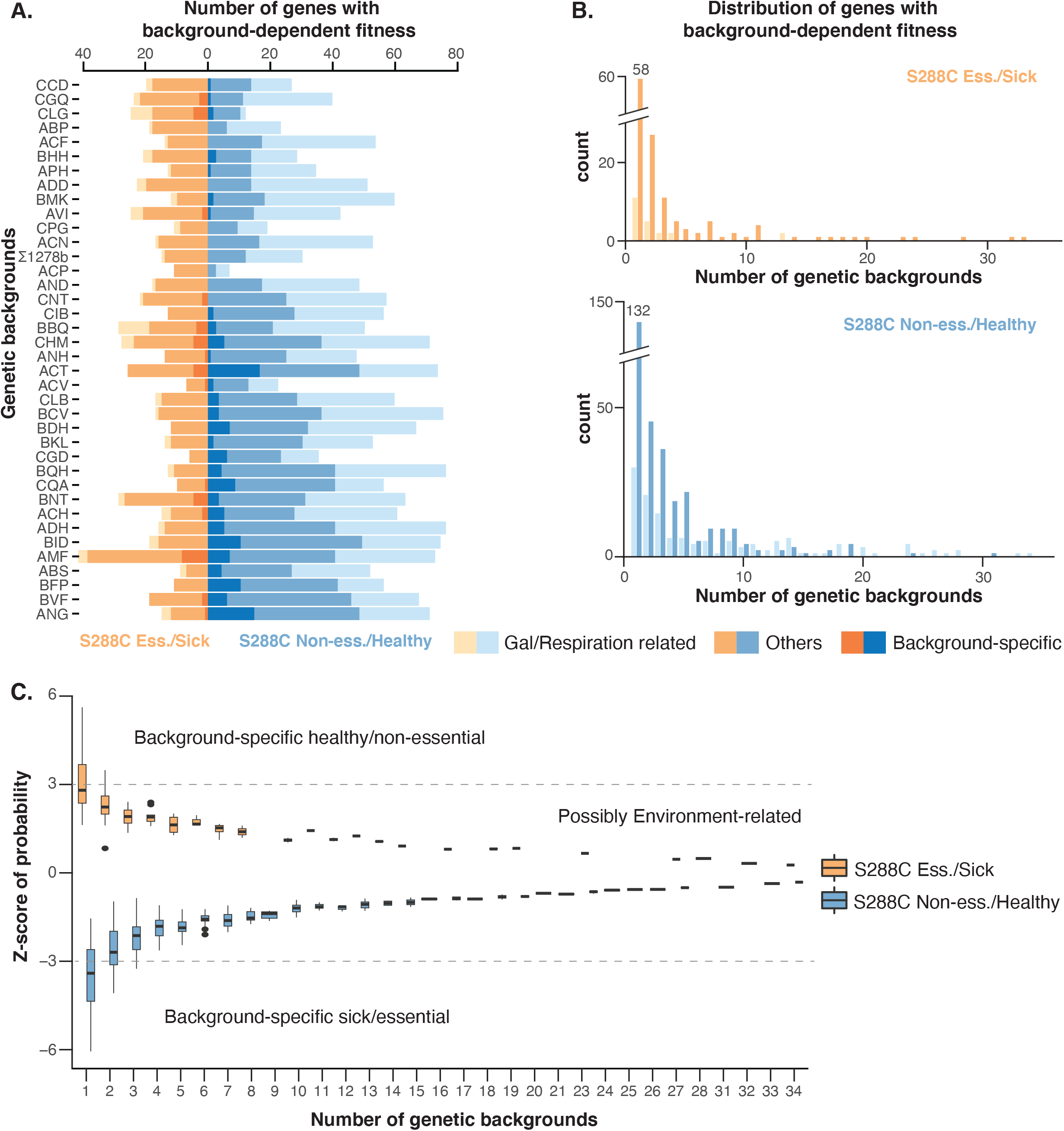
Number and distribution of background-dependent fitness variation genes. (A) Number of hits detected in each genetic background. Genes annotated as galactose or respiration-related and background-specific genes are color-coded as indicated. Strains are sorted according to the total number of insertions. (B) The number of genetic backgrounds impacted by the detected hits. Top panel, gain-of-fitness genes compared to S288C; bottom panel, loss-of-fitness genes compared to S288C. (C) Z-statistic distribution for hits that impact different numbers of genetic backgrounds. A cut-off of |z-statistics| > 3 is indicated with dotted lines.

We then calculated the z-statistics for all variable fitness hits to distinguish those that are background-specific from the others that are possibly related to the environment (Table S4). In principle, environment-related cases are more likely to vary continuously in the population with a low z-statistics, whereas cases that are truly specific to some genetic backgrounds should be outliers with a high z-statistic score (|z| > 3) (Figure 3C). Of the set of 632 genes, we found 179 that are background specific, which mainly impact a single genetic background (Table S4). These background-specific genes are rarer compared to the environment-related group, with a median of 5 identified per isolate both loss- and gain-of-fitness types combined (Figure 3A). Genes related to respiration and mitochondrial functions are not overrepresented in this group (23/179 *vs*. 691/4469 in the background, Fischer’s exact test P-value = 0.82). No significant enrichment for any biological processes or molecular functions has been identified. By contrast, respiration-related genes are significantly overrepresented in the remaining group (140/453 *vs*. 691/4469, Fischer’s exact test P-value = 1.6e-10, odds ratio = 2). Each of these 453 potentially environment-related genes impact on average 6 genetic backgrounds.

### Environment-related fitness variation reveals potential functional rewiring

While a large fraction of potentially environment-related hits correspond to genes known to be involved in respiration, the majority of this group is involved in other biological processes. To explore the functional relationships within this group, we calculated the pairwise correlations between these genes using predicted fitness values across the 39 strain backgrounds (Figure 4A). We constructed a network based on the profile similarities where the edges correspond to a Pearson’s correlation > 0.6 (correlation) or < -0.6 (anti-correlation) (Figure 4B, Figure S4). In total, 292 out of the 453 environment-related hits exceeded our stringent correlation cut-offs (Figure S4). The profile similarity and network structure revealed two main subnetworks, which are correlated within the subgroup but are anti-correlated between subgroups (Figure 4A-B). The first subgroup contains mainly respiration-related genes, in particular genes involved in mitochondrial translation (Figure 4A, Figure S4A), which are anti-correlated with genes involved in transcription regulation and chromatin remodeling (*SPT7, SPT8, SWC4, SWC5, ARP6, ARP7, SIN3, RKR1, YAF9, UME1, NGG1, CHD1, STH1*, for example) as well as genes involved in nuclear-cytoplasmic protein transfer (*KAP120, KAP122, KAP123, NUP57, NUP100, NUP188, POM152, NIC96, MLP1*, for example) (Figure S4A). Many of these correlations have been found between members of the same protein complexes. Several members of the transcription and nuclear transport subgroup are also annotated as related to respiration (deletion leads to absence of respiration) although they are not directly involved in mitochondrial function, such as *SIN3*, a general chromatin remodeler, and *KAP123*, a karyopherin responsible for nuclear import of ribosomal proteins. In addition to this large network, several small networks have also been detected (Figure S4B-D), including *PMT1, PMT2* and *GET2*, which are involved in ER related glycosylation and are known to have physical interactions. Functional enrichments in the anti-correlated subgroups suggest a potential ‘rewire’ between mitochondrial translation and transcription regulation/nuclear transport, where modular switched of fitness effects associated with gene loss-of-function may occur in different genetic backgrounds.

**Figure 4.**
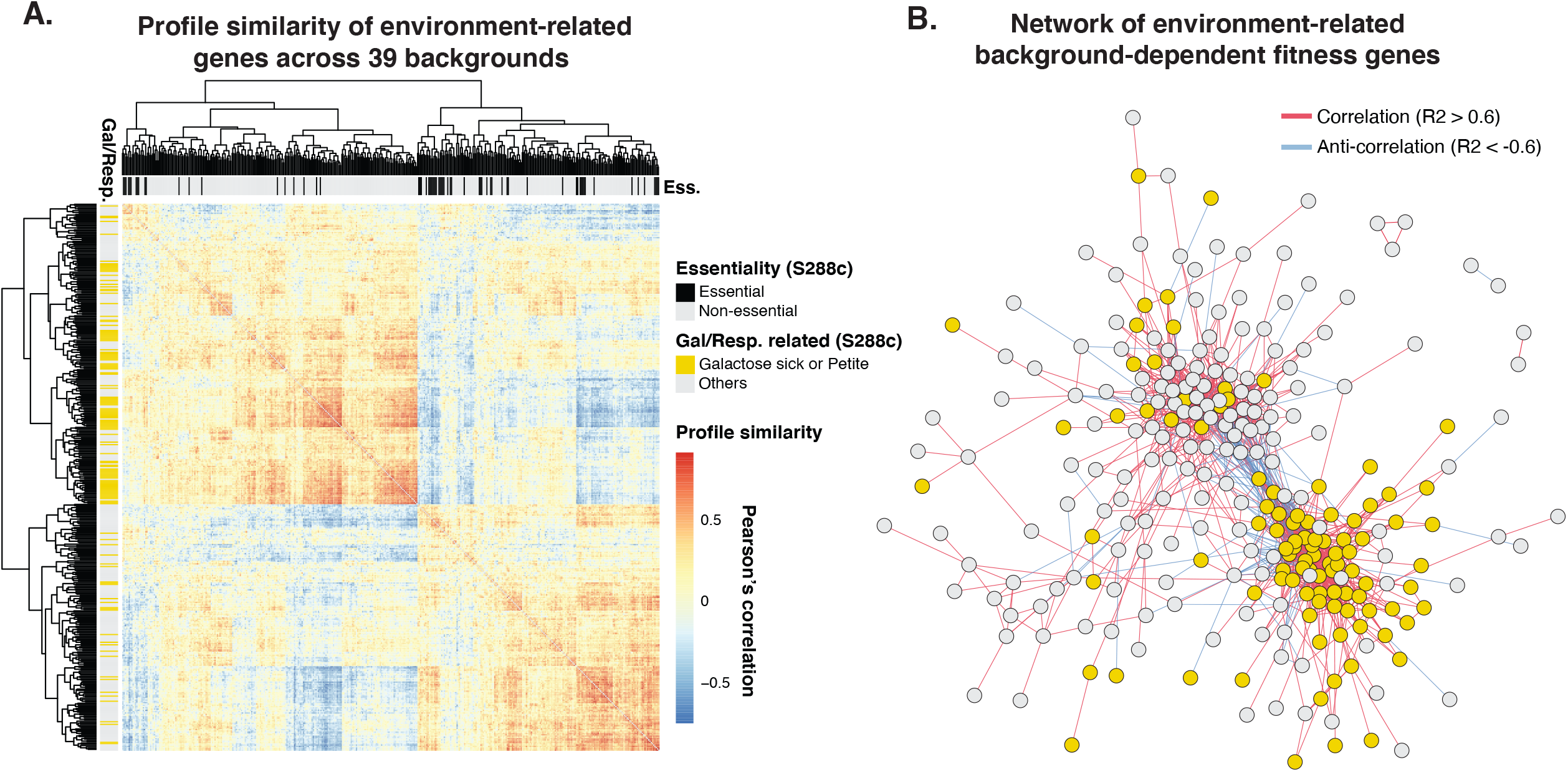
Correlation analyses for environment-related hits. (A) Pairwise profile similarity based on predicted fitness across 39 backgrounds. Distance matrix was based on pairwise Pearson’s correlation. Gene essentiality annotations are indicated on the upper sidebar and genes annotated as involved in galactose/respiration are indicated on the left sidebar. (B) Network based on profile similarity among environment-related hits. Genes annotated as involved in galactose/respiration are colored in yellow. Positive correlations (> 0.6) are represented as red edges and negative correlations (< -0.6) are represented as blue edges. Complete network with annotated gene names can be found in Figure S4.

### Functional insights into background dependent fitness genes

To further explore the functional enrichments of fitness variation genes at the strain level, we annotated genes in our dataset into 16 functional neighbourhoods according to SAFE^37^ and looked for enrichment in different neighbourhoods (Figure 5A). For each neighbourhood, we calculated the odds ratio of enrichment based on the number of hits annotated in the neighbourhood *vs*. the total number of hits, with the size of the neighbourhood and the total number of genes as background (one-sided Fisher’s exact test). Globally, background-specific hits are not enriched for most processes except for cell polarity (OR = 1.49, P-value = 0.026). Environment-related hits are enriched for respiration and mitochondrial functions (OR = 3.77, P-value = 4.16e-17), as well as transcription and chromatin regulation (OR = 1.53, P-value = 0.002), nuclear cytoplasmic transport (OR = 2.14, P-value = 0.004) and DNA repair (OR = 1.55, P-value = 0.01) (Figure 5A, Table S4). When looking at the same neighbourhood enrichment at the strain level, environment-related hits are enriched for mitochondrial functions in most genetic backgrounds, with the exception of ACP and CLG strains, the latter of which has a predicted fitness profile that was most similar to the reference S288C (Figure 2). A large fraction of isolates showed significant enrichments for transcription and chromatin regulation as well as nuclear-cytoplasmic transport (Figure 5A). These enrichments are consistent with the rewiring hypothesis based on the profile similarity network analysis (Figure 4B). Indeed, by specifically looking at the annotated genes in these functional neighbourhoods, we observed various degrees of rewiring depending on the backgrounds (Figure 5B-C). In the reference S288C, loss-of-function for annotated genes in these three neighbourhoods showed either high- or low-fitness predictions (Figure 5B, Figure S5A). While in other genetic backgrounds, these predictions may be reversed as gain- or loss-of-fitness hits compared to S288C, with profiles ranging from similar to S288C (CLG) to almost completely reversed (AMF) (Figure 5C). Most notably, such rewire could include either only mitochondrial-related genes, or with one or more processes related to either transcription and chromatin regulation or nuclear-cytoplasmic transport (Figure 5C). Depending on the genetic background, different sets of genes within the same functional neighbourhood could be involved, highlighting the dynamics of such rewire (Figure S5B).

**Figure 5.**
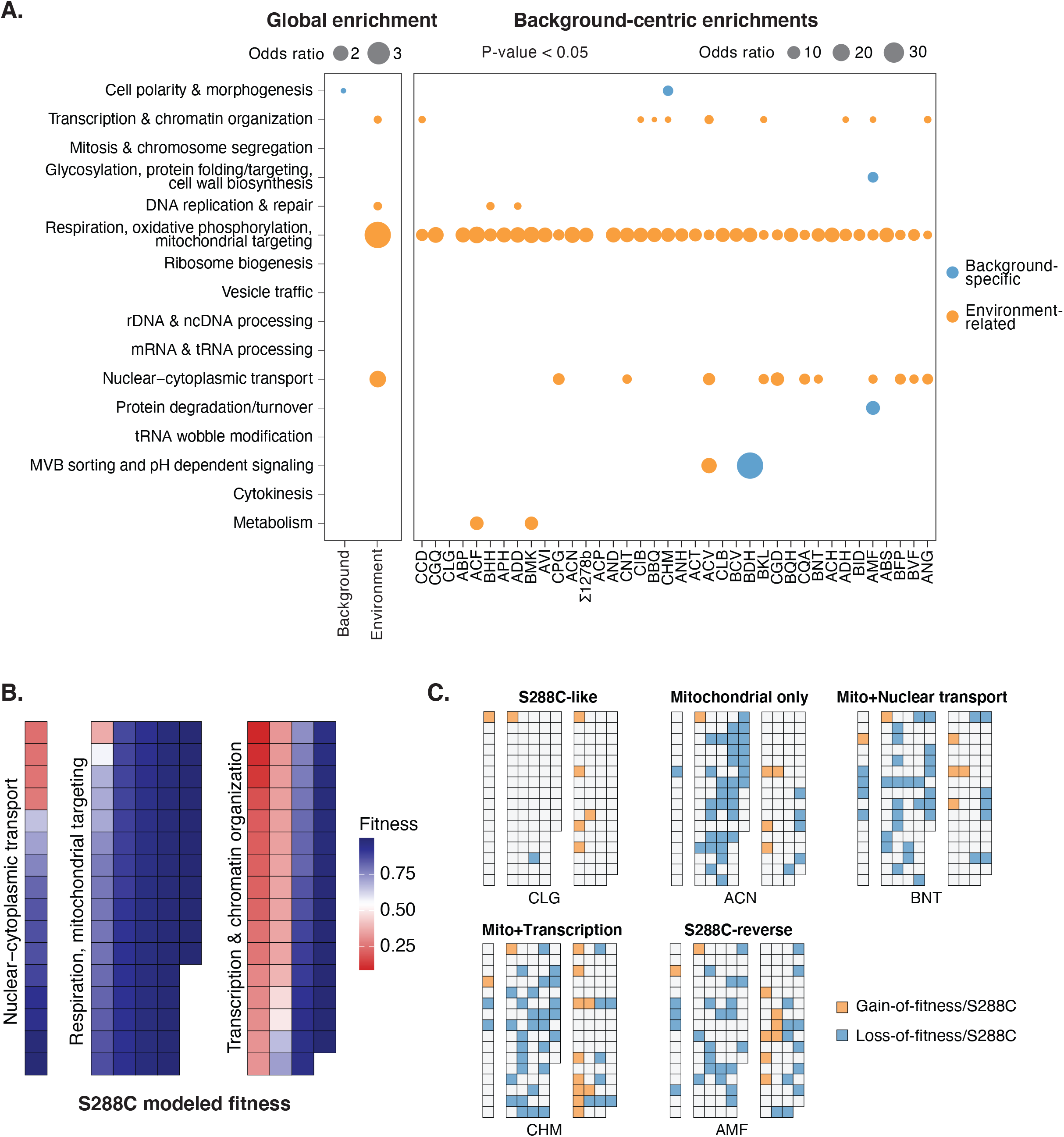
Functional enrichments and rewiring for background-dependent fitness genes. (A) Enrichments across 16 functional neighbourhoods defined by SAFE^37^. Dot sizes represent odds ratios between the number of hits in a given neighbourhood *vs*. the total number of hits detected, with the size of the neighbourhood *vs*. the total number of genes in the dataset as background, using one-sided Fischer’s exact test. Global enrichment for background-specific (blue) and environment-related (orange) hits are presented on the left panel, and strain-centric enrichments are on the right panel. Enrichments with a p-value < 0.05 are shown. Backgrounds highlighted by dashed lines corresponds to example rewiring diagrams in (C). (B) Predicted fitness for genes annotated in Respiration/mitochondrial targeting, Transcription and chromatin organization and nuclear-cytoplasmic transport in the reference S288C. Genes in different processes are arranged by descending order of the modeled fitness. Detailed annotated version of this diagram can be found in Figure S5A. (C) Example rewiring diagrams in other backgrounds compared to the reference S288C. A switch from healthy to sick (loss-of-fitness) is indicated in blue and a switch from sick to healthy (gain-of-fitness) is indicated in orange for any given gene in a given background. The diagrams for all 38 isolates are shown in Figure S5B.

Compared to environment-related genes, background-specific ones are rare, and tend to show little functional enrichment, as expected. However, in cases where multiple genes are detected in the same genetic background, some enrichments emerge (Figure 5A, Table S4). For example, in the strain BDH, 8 background-specific genes were detected with 3 annotated into one of the 16 functional neighbourhoods, and two of which are involved in MVB sorting and pH dependent signaling (*RIM8 & RIM101*). Both genes are non-essential in S288C but predicted as loss-of-fitness in the BDH background (Figure 5A). In the strain AMF, 16 background-specific genes were detected with 11 annotated, among which 2 were involved in protein degradation and turnover (*VID28 & PRE3*) and 3 were involved in glycosylation and cell wall biogenesis (*OST1, OPI3 & FAB1*). These observations demonstrate that background-specific fitness variation genes, while rare, can be functionally related and may involve multiple members of the same protein complex or biological process.

Finally, as previously posited^6^, genes exhibiting background-dependent fitness variation tend to show an intermediate level of connectivity in terms of genetic interactions (Figure 6A, Table S5) and an intermediate functional similarity between interacting gene pairs compared to genes that are consistently non-essential or essential (Figure 6B). Both environment-related and background-specific hits have the same pattern. Interestingly, background-specific genes display higher non-synonymous to synonymous substitution rates (dN/dS) than essential genes and non-essential genes (Figure 6C), indicating potential positive or relaxed purifying selection on these genes at the population level. Overall, background-specific fitness genes tend to be diverse yet can be functionally related within a single genetic background. Genes with environment-related fitness variation share general evolutionary characteristics with background-specific cases.

**Figure 6.**
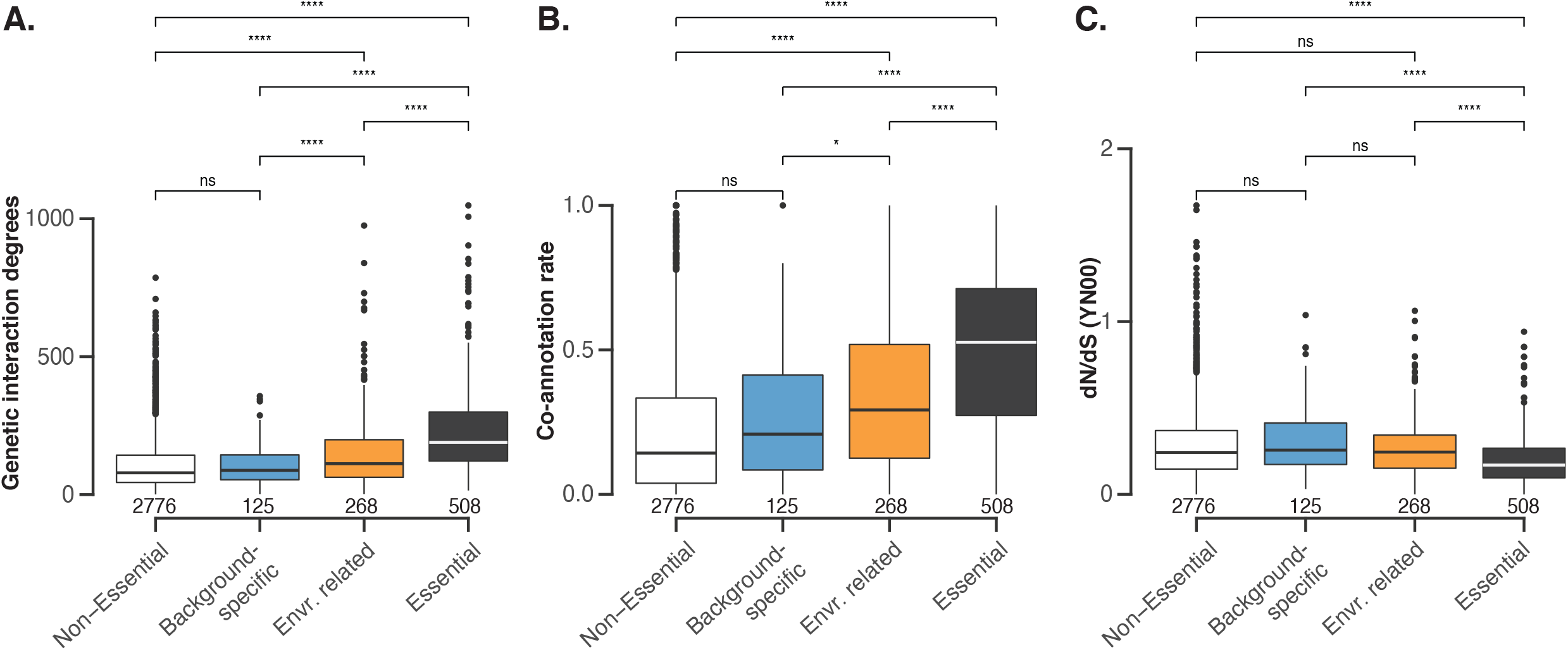
Evolutionary features associated with background-dependent fitness genes. (A) Genetic interaction degrees derived from the yeast global genetic interaction network^37^ for non-essential, background-specific, environment-related and essential gene categories. The number of genes annotated in each category are indicated. (B) Functional co-annotation rates^37^ for different gene categories. The co-annotation rate corresponds to the fraction of interaction partners that are annotated in the same biological process as the primary gene^37^. (C) Mean non-synonymous *vs*. synonymous substitution rates (dN/dS) across 1,011 natural yeast isolates using the YN00 method^26^. Comparisons between categories were performed using T-test, and significance levels are as indicated, with ns: P-value > 0.05, *: P-value < 0.05, **: P-value < 0.01, ***: P-value < 0.001 and ****: P-value ≪ 0.0001.

## Discussion

To have a better insight into the background-dependent fitness variation associated with gene loss-of-functions, we explored a large number of natural yeast isolates using a transposon saturation strategy. We modeled fitness by considering transposon insertion densities within gene coding sequence and surrounding regions. The comparison of the modeled fitness between different isolates and the reference S288C allowed the identification of 632 genes displaying background-dependent phenotypes. The majority of these cases (71,7%) showed continuous fitness variation across the population and is at least partly related to the environment. By contrast, background-specific cases tend to be rare, with on average 5 genes per isolate. At the individual level, both environment-related and background-specific variable fitness genes can be functionally related.

The impact of the environment on the background-dependent fitness genes can be supported by two main observations. First, this set of genes was highly enriched for respiration and mitochondrial functions, which is consistent with a fitness loss under prolonged growth in media with galactose as the sole carbon source. Indeed, mitochondrial-related genes were also found to be background-dependent in a previous study involving 4 different isolates under conditions with non-fermentable carbon sources^25^. Second, these genes showed a continuous variation across the population. Further analyses highlighted that genes involved in two biological processes, namely transcription & chromatin remodeling and nuclear-cytoplasmic transport, are anticorrelated with genes involved in mitochondrial translation in terms of their fitness profiles. These anticorrelations indicate a modular change in the relative fitness of genes involved in these processes compared to the reference strain S288C. However, whether such rewiring effect is exclusively related to respiration conditions or could represent a general background-dependency effect remains difficult to disentangle due to the experimental conditions required for transposon saturation analyses.

In a recent large-scale analysis of environment-dependent genetic interactions, it has been shown that most interactions specific to an environmental condition are in fact part of the global genetic interaction network that were exacerbated or attenuated in the tested condition^38^. Compared to genetic interactions between pairs of gene deletion mutants, the background-dependent gene loss-of-function phenotype could be considered as interactions between the loss-of-function gene and background-specific modifiers, which are expected to share general properties to genetic interactions with deletion mutants. Indeed, we tested the gene deletion phenotype for one of the environment-related genes involved in transcription and chromatin remodeling, the *BMH1*gene (Figure S5C). This gene was identified as loss-of-fitness in multiple genetic backgrounds compared to S288C in our study. Interestingly, the loss-of-fitness phenotype was indeed confirmed on standard rich media, suggesting the environment-related fitness variation genes could have a general effect. In addition, genes involved in chromatin remodeling were also found to display background-dependent fitness effects in a previous study comparing S288C and a natural isolate 3S^5^. These observations suggest that the rewiring effect could have implications beyond a specific experimental condition.

Although transposon saturation strategy can be versatile to genetic diversity among isolates, this method also presents some limitations. Among all the isolates initially tested, only about half showed a reasonable level of insertion efficiency, highlighting the unexpected variability of transposon activity between different individuals. This variability results in an underestimate of the number of genes associated with background-dependent phenotypes. In addition, loss-of-function phenotypes that are related to specific protein domains but not the entire ORF are difficult to identify, unless the insertion efficiency is extremely high. The *Hermes* system, like all currently available saturation systems, requires step of a transposon induction in the presence of galactose^30^. This competition effect in a non-fermentable carbon source may complicate downstream analysis as the effects of environment *vs*. genetic background can be difficult to unravel. New strategies that take into account these factors are still needed in order to get a more precise view of background-dependent gene loss-of-function phenotypes at the species level.

## Material and methods

### Strains and growth conditions

A total of 106 isolates were selected from the 1,011 *Saccharomyces cerevisiae* collection^26^. A prototrophic haploid strain FY5, isogenic to the reference strain S288C was also included. Haploid segregants derived from the 106 natural isolates were obtained after *HO* deletion and tetrad dissection^32,33^. Detailed descriptions of the strains can be found in Table S1. Strains were maintained at 30°C using YPD (1% Yeast extract; 2% Peptone, 2% Dextrose) in liquid culture or solid plates (2% of agar). Transposon activity was induced in YPGal (1% Yeast extract; 2% Peptone, 2% Galactose) with Hygromycin B (200 µg/mL). Sporulation was induced on solid plates containing 1% of potassium acetate and 2% of agar.

### Ploidy control

Ploidy was estimated by flow cytometry. Cells in exponential growth phase were washed in water, then 70% ethanol and sodium-citrate buffer (50 mM, pH 7.5) followed by RNase A treatment (500 µg/mL). To avoid cell aggregates, each sample was sonicated then the DNA was labelled with propidium iodide (16 µg/mL), a fluorescent intercalating agent. DNA content was then quantified using the 488 nm excitation laser of the Accuri C6 plus flow cytometer (BD Biosciences).

### Cell transformation

Cells in exponential growth phase were chemically transformed using the EZ-Yeast Transformation Kit (MP biomedicals). We incubated cells 30 minutes at 42°C with EZ-Transformation solution, carrier DNA and either 100 ng of pSTHyg plasmid or 1 µg of PCR fragment. After regeneration in YPD, cells were spread on solid YPD plate supplemented with Hygromycin B and incubated at 30°C until transformants appeared.

### Construction of the pSTHyg plasmid

In order to be compatible with our isolates already carrying either a nourseothricin or a kanamycin resistance cassette, the nourseothricin cassette of the pSG36 plasmid^29^ was replaced by a hygromycin B resistance cassette. The pSG36 plasmid was amplified in 2 fragments by PCR excluding the *natMX* cassette, then assembled with the *hphMX* cassette amplified from p41 plasmid (Addgene #58547) with overlapping regions using Gibson assembly. The new plasmid, pSTHyg was amplified in *E. coli* and extracted using the GeneJET Plasmid Miniprep Kit (Thermo Scientific™). The construction was verified using enzymatic digestion with *KpnI* and *PvuI*.

### Generation of transposon insertion mutant pools

Each natural isolate was grown in liquid YPD medium and chemically transformed with 100 ng of pSTHyg plasmid as described. From the selective transformation plates, a single clone was picked and grown in 30 ml of YPD supplemented in hygromycin B under agitation at 30°C until saturation (∼ 24h). Cells were then diluted at an OD of 0.05 in 50 ml of YPGal supplemented with hygromycin B to activate the transposase and induce the transposition for 72h at 30°C. Two successive dilutions were then performed for 24h at an OD of 0.5 in 100 ml of YPD then YPD supplemented with hygromycin B to enrich for cells the transposon in their genome. The final 100ml culture was centrifuged, water-washed and 500 µl aliquots of cells were frozen at -20°C.

### Sequencing library preparation

In order to sequence the genomic regions with a transposon insertion, the genomic DNA of the pool of cells carrying insertion events was extracted using the MasterPure™ Yeast DNA Purification Kit (Lucigen). Cells were lysed using a lysis solution supplemented in zymolyase 20T (1.5 mg/ml). Proteins and cellular debris were removed with the MPC Protein Precipitation Reagent and several RNase A treatments were realized to eliminate RNA. Genomic DNA was then precipitate with ethanol. The pellet was washed twice with 70% ethanol and resuspended in 80 µl of water. The gDNA sample integrity was controlled on 1% agarose gel and quantified on Nanodrop and Qubit using the Qubit™ dsDNA BR Assay Kit (Invitrogen™). 2 × 2 µg of gDNA were digested in parallel with 50 units of *DpnII* (NEB #R0543L) and *NlaIII* (NEB #R0125L) in 50 µl for 16h at 37°C. The enzymatic reactions were inactivated for 20 min at 65°C and DNA fragments were ligated with 25 Weiss units of T4 Ligase (Thermo Scientific #EL0011) in a total volume of 400 µl for 6h at 22°C. Circular DNA were then precipitated overnight at -20°C with ethanol, salt (NaOAc 3M pH5.2) and glycogen. After an 70% ethanol wash, the DNA pellet was resuspended in 50µL of water. The junction between the genomic region and the transposon insertion site was amplified on both *DpnII* and *NlaIII* digested and re-circularized gDNA by PCR using outward-facing primers targeting the transposon. The PCR products were controlled on 1% agarose gel and displayed variable sizes centred around 750 bp. Nanodrop and Qubit using the Qubit™ dsDNA BR Assay Kit (Invitrogen™) quantifications were then performed to pool the same amount of *NlaIII*-digested and *DpnII*-digested PCR products. For each sample, at least 6 µg at minimum 30 ng/µl was then sent to the BGI (Beijing Genomics Institute) for sequencing. In total, each sequencing run provided 1 Gb of 100 bp paired-end reads using Illumina Hi-Seq 4000 or DNBseq technologies.

### Determination of transposon insertion sites

The reads that contained the amplified part of the transposon were selected and the corresponding 57 bp sequence was trimmed with Cutadapt^39^ and the reads corresponding to the plasmid were discarded. The cleaned reads were mapped to the S288C reference genome with the corresponding SNPs inferred for each isolate^26^ with BWA^40^. The genomic position of an insertion site was defined as the first base pair aligned on the genome after the transposon region. For each insertion site, the number of reads and their orientation were obtained.

### Modelling the fitness effect of gene loss-of-function based on transposon insertion profiles

The number of insertions in the promoter region (−100 bp to ATG), beginning of the coding region (−100 bp to +100 bp from ATG), the coding region, end of the coding region (−100 bp to +100 bp from stop-codon) were normalized as insertion densities per 100 bp. Gene essentiality annotations were obtained from SGD (phenotype “inviable”) exclusively for annotations with gene deletion in the S288C background. Respiration related gene annotations were obtained from SGD with the phenotype “respiration: absent” after gene deletion in S288C. Galactose-specific loss-of-fitness was determined in Costanzo et al.^38^, with a stringent cut-off of < -0.2. A logistic model was constructed using the glm() function from the R package “stats”, using insertion densities in the reference strain S288C, in the promoter region (−100 bp to ATG), beginning of the coding region (−100 bp to +100 bp from ATG), the coding region, end of the coding region (−100 bp to +100 bp from stop-codon), raw insertion number in the coding region and gene sizes as predictors, and essentiality annotations as a binary classifier. Genes displaying a slow growth phenotype^41^, genes with differential fitness defect in galactose media^38^, as well as genes showing respiration defects were excluded. Genes that are localized in regions with low insertion densities, *i*.*e*. less than 3 insertions in the terminator region (STOP to +300 bp) and less than 50 insertions in a 10 kb region surrounding the gene (−5 kb before ATG and +5 kb after STOP) were also excluded. A total of 4600 genes were included in the model corresponding to 867 essential genes and 3737 non-essential genes (Table S2). 10-fold cross-validation was performed using the R package “caret”, with trainControl() and train() functions, method = “glm”, family = “binomial”. Cross-validation results showed an average accuracy of 0.88 with a Kappa 0.57 (Table S2). The predictive value for non-essential labels is 0.91, contrasting to a lower predictive value of 0.70 for essential labels, indicating a better accuracy in predicting non-essential genes using this model. This lower predictive power for essential genes is more or less expected as the absence or low numbers of insertions could be linked to the overall low insertion density in certain genomic regions, which is independent of gene essentiality. Imputations for missing values were performed using the function impute.knn() in the R package “impute”, with k = 10, rowmax = 50% and colmax = 80%. Quantile normalization of the imputed matrix was performed using normalize.quantiles() function in the R package “preprocessCore”. All fitness prediction data can be found in Table S3.

### Validation of the phenotypic consequence of *BMH1* gene loss-of-function

Stable haploid isolates, FY5 and CIB were diploidized using the pHS2 plasmid (Addgene #81037) containing the *HO* gene encoding the endonuclease responsible for mating type switching and a hygromycin resistance cassette. The *BMH1* gene was replaced with a Hygromycin B resistance cassette in the diploid isolates. Sporulation was induced on potassium acetate medium in diploid isolates heterozygous for *BMH1* gene deletion. Around 20 resulting tetrads were then dissected on YPD using a MSM 400 micromanipulator (Singer Instrument). Each spore grew for 48h at 30°C and the colony size was captured with the camera of the colony picker, PIXL (Singer Instrument). Colony size measurements were then analysed using custom R scripts.

## Data availability

All sequencing data related to this study were deposited to the European Nucleotide Archive (ENA) under the accession number PRJEB45777.

## Acknowledgments

We thank Agnès Michel for helpful suggestions throughout the project. This work was supported by the European Research Council (ERC Consolidator Grant 772505). It is also part of the Interdisciplinary Thematic Institute IMCBio, as part of the ITI 2021-2028 program of the University of Strasbourg, CNRS and Inserm, supported by IdEx Unistra (ANR-10-IDEX-0002), and EUR (IMCBio ANR-18-EUR-0016) under the framework of the French Investments for the Future Program. JS is a member of the Institut Universitaire de France.

## Supplemental figure legends

**Figure S1.**
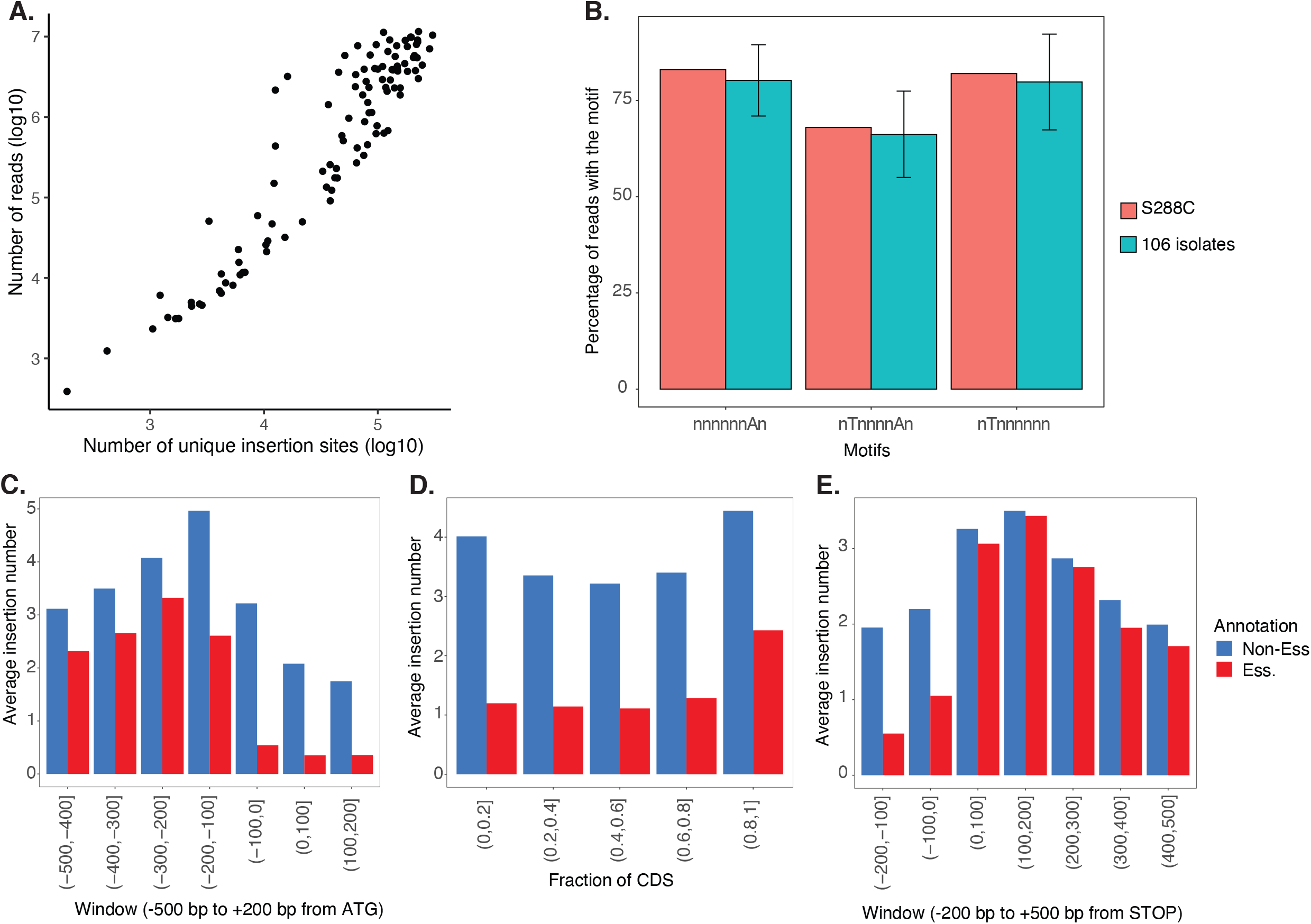
(A) Number of reads (y-axis, log10 scale) *vs*. number of unique insertion sites (x-axis, log10 scale) across 107 diverse isolates. (B) Insertion preference comparison between the reference S288C and the other 106 selected isolates. Sequence motifs are on the x-axis and the percentage of reads with a given motif are presented as color coded bars. Error-bars correspond to the standard deviation across different isolates. (C) Insertion density comparison between essential and non-essential genes in S288C in the promoter region. Average insertion numbers in the -500 bp to +200 bp region relative to ATG are shown in 100 bp windows. (D) Insertion density comparison between essential and non-essential genes in S288C in the coding region (CDS). Average insertion numbers in the relative fractions of a given CDS are shown. (E) Insertion density comparison between essential and non-essential genes in S288C in the terminator region. Average insertion numbers in the -200 bp to +500 bp region relative to the stop-codon are shown in 100 bp windows.

**Figure S2.**
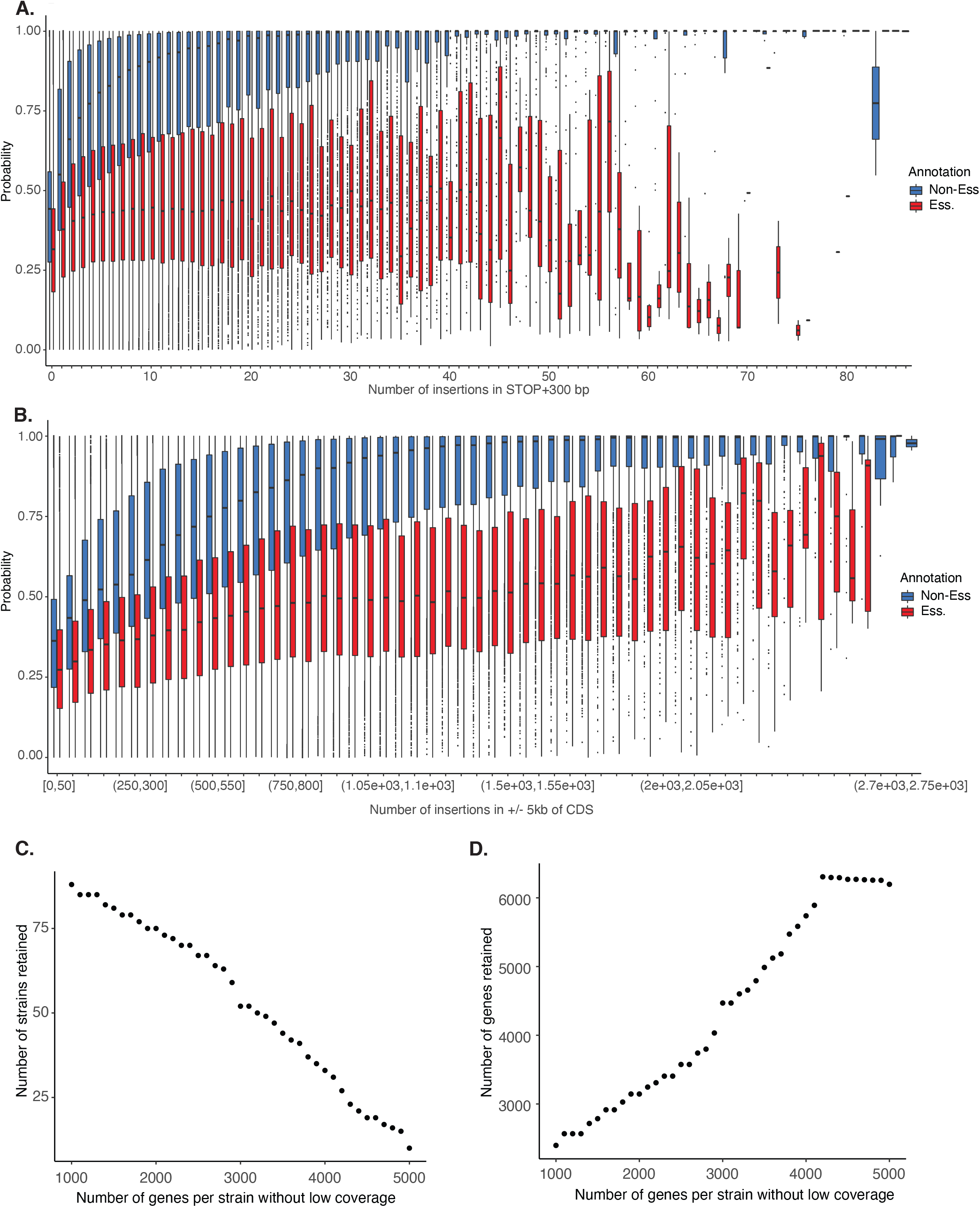
(A) Predicted non-essential probabilities (y-axis) as a function of the number of insertions in the terminator region (300 bp after stop-codon). Non-essential genes are in blue and essential genes in red. (B) Predicted non-essential probabilities (y-axis) as a function of the number of insertions in a 10 kb region surrounding the CDS (5 kb before ATG and 5 kb after stop-codon). Non-essential genes are in blue and essential genes in red. (C) The number of strains retained as a function of cut-offs of the number of interpretable genes after removing low coverage regions (less than 50 insertions in the surrounding 10kb region and/or less than 3 insertions in the 300 bp terminator region). (D) Number of genes retained after imputation as a function of cut-offs of the number of interpretable genes after removing low coverage regions.

**Figure S3.**
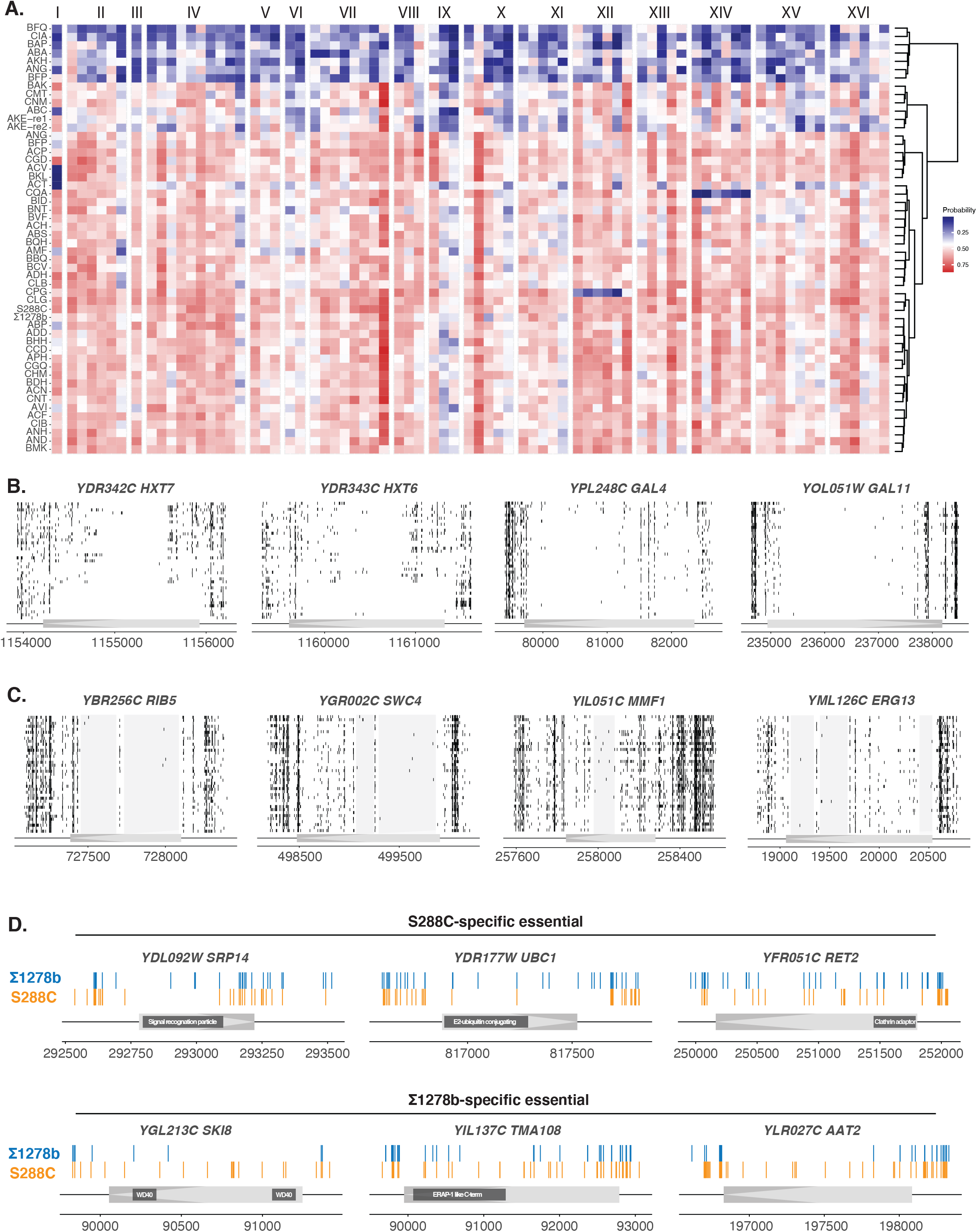
(A) Average non-essential probability or predicted fitness for every 10 successive essential genes along all 16 chromosomes for 52 strains that passed the coverage cut-offs. Strain-side clustering was based on predicted fitness for all genes. (B) Insertion profiles for gene related to galactose metabolism that are annotated as non-essential in S288C but detected as essential/sick in all or a fraction of the 39 strains in the final dataset. Chromosomal positions and gene orientations are schematically presented on the x-axis and insertion profiles for each strain are presented as black vertical bars. (C) Insertion profiles for essential genes predicted as non-essential in S288C. Shaded areas correspond to potential essential protein domains. (D) Insertion profiles for genes previously shown background-specific essentiality between S288C and Σ1278b. Domain-specific essentiality regions are indicated.

**Figure S4.**
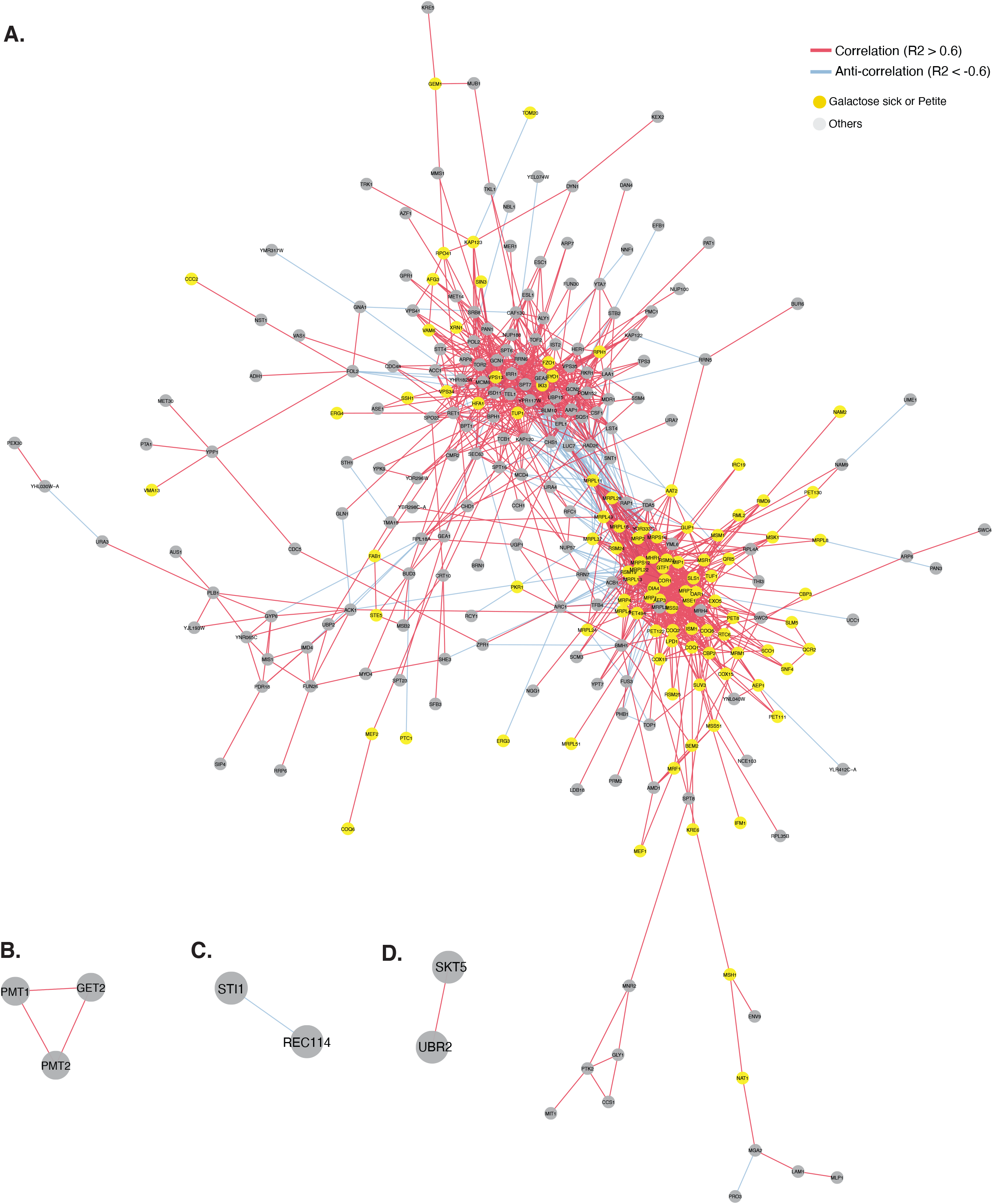
(A) Annotated network based on profile similarity as shown in Figure 4B. (B-D) Subnetworks with significant correlations independent from the large subnetwork involving respiration-related hits.

**Figure S5.**
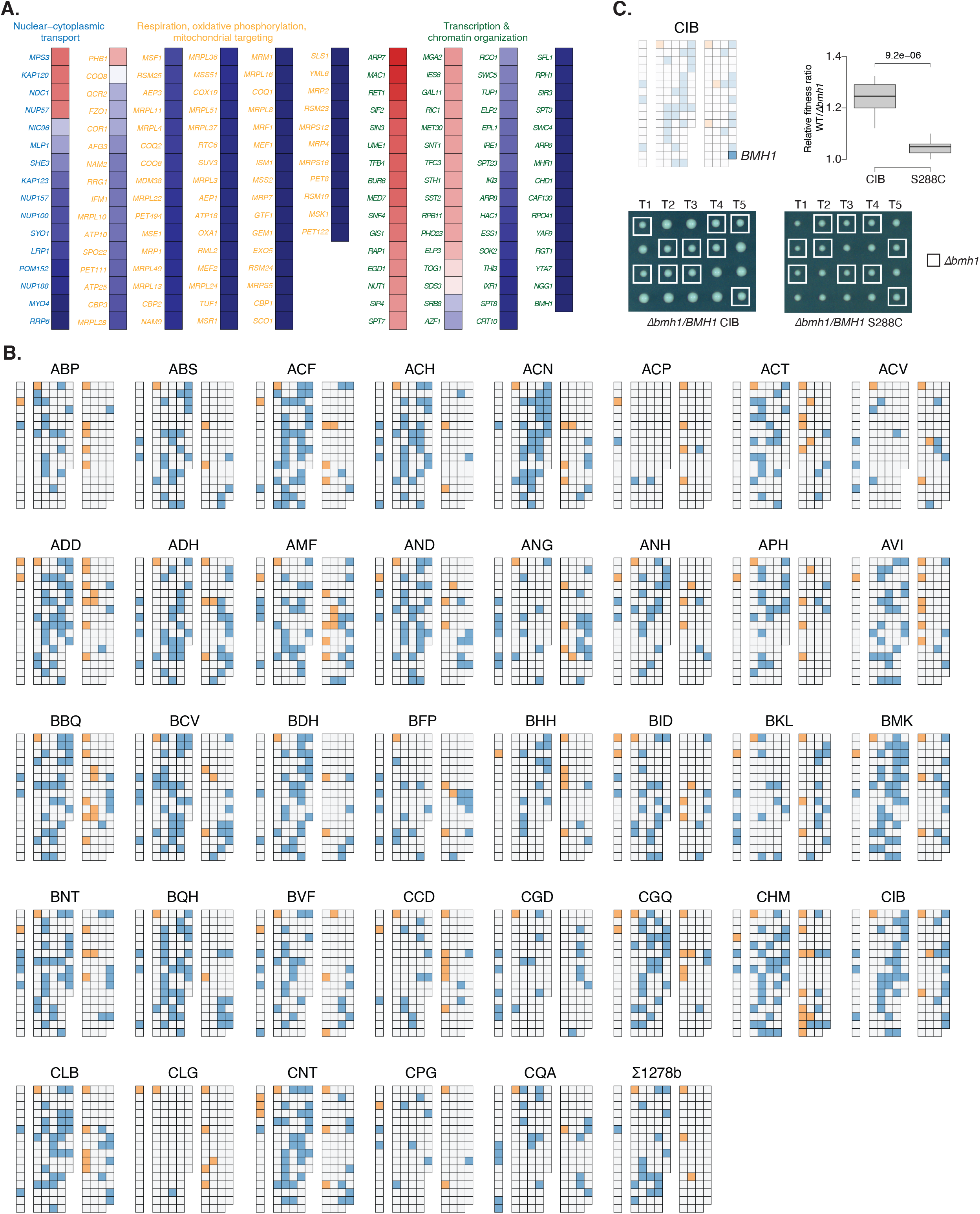
(A) Predicted fitness for genes annotated in respiration/mitochondrial targeting, transcription & chromatin organization and nuclear-cytoplasmic transport in the reference S288C with gene name annotations. Related to Figure 5B. (B) Rewiring diagrams for all 38 isolates relative to the reference S288C. Related to Figure 5C. (C) Example of functional rewire in a natural isolate CIB compared to the reference S288C for a transcription-related hit *BMH1*. Relative fitness ratio (colony size for WT divided by deletion of *BMH1*) is shown on the upper right panel. Colony sizes of *BMH1* deletion *vs*. WT were measured using tetrad dissection of hemizygous diploids. 5 tetrads are shown for each background.

## Supplemental tables

**TableS1- Description of isolates used in this study**

**TableS2- Model construction and evaluations**. This table contains 4 tabs:

GenesInModel: 4600 ORFs and their essentiality annotations used to construct the logistic model. Insertion numbers and densities within coding sequence and surrounding regions are included. Insertion numbers calculated from S288C insertion profile.

ModelSummary: Features included in the logistic model and their coefficient.

CrossValidation: Summary of the cross validation results.

CMStat: Confusion matrix, prediction accuracy and precision for essential/non-essential labels.

**TableS3- Raw and final dataset with predicted fitness**. This table contains 3 tabs:

Raw_data_pred: All raw predicted fitness based on the logistic model for 107 isolates.

Pred_final_39: Predicted fitness for 39 isolates included in the final dataset. Raw, imputed and quantile normalized predictions are shown.

Score_final_39: Differential fitness score by comparing the predicted fitness in a given isolate to S288C.

**TableS4- Background-dependent fitness variation genes identified in this study**. This table contains 4 tabs:

Z-statistics: Z-statistics for each of the 632 hits, including the number of genetic backgrounds impacted for each hit.

Hits_SAFE_annotation: Annotations for each hit into the 16 functional neighbourhoods according to SAFE^37^.

Enrichment_global: Enrichment for all hits across 16 functional neighbourhoods

Enrichment_Strain: Enrichment for hits in a given genetic background across 16 functional neighbourhoods.

**TableS5- Genetic interaction degree and dN/dS values and gene classifications**

## References

1. Cooper, D. N., Krawczak, M., Polychronakos, C., Tyler-Smith, C. & Kehrer-Sawatzki, H. Where genotype is not predictive of phenotype: towards an understanding of the molecular basis of reduced penetrance in human inherited disease. Hum. Genet. 132, 1077–1130 (2013).

2. Chandler, C. H., Chari, S., Tack, D. & Dworkin, I. Causes and Consequences of Genetic Background Effects Illuminated by Integrative Genomic Analysis. Genetics 196, 1321–1336 (2014).

3. Sackton, T. B. & Hartl, D. L. Genotypic Context and Epistasis in Individuals and Populations. Cell 166, 279–287 (2016).

4. Chow, C. Y. Bringing genetic background into focus. Nat. Rev. Genet. 17, 63–64 (2016).

5. Mullis, M. N., Matsui, T., Schell, R., Foree, R. & Ehrenreich, I. M. The complex underpinnings of genetic background effects. Nat. Commun. 9, 3548 (2018).

6. Hou, J., van Leeuwen, J., Andrews, B. J. & Boone, C. Genetic Network Complexity Shapes Background-Dependent Phenotypic Expression. Trends Genet. 34, 578–586 (2018).

7. Chen, R. et al. Analysis of 589,306 genomes identifies individuals resilient to severe Mendelian childhood diseases. Nat. Biotechnol. 34, 531–538 (2016).

8. Fournier, T. & Schacherer, J. Genetic backgrounds and hidden trait complexity in natural populations. Curr. Opin. Genet. Dev. 47, 48–53 (2017).

9. Chow, C. Y., Kelsey, K. J. P., Wolfner, M. F. & Clark, A. G. Candidate genetic modifiers of retinitis pigmentosa identified by exploiting natural variation in Drosophila. Hum. Mol. Genet. 25, 651–659 (2016).

10. Steinberg, M. H. & Sebastiani, P. Genetic modifiers of sickle cell disease. Am. J. Hematol. 87, 795–803 (2012).

11. Dorfman, R. Modifier gene studies to identify new therapeutic targets in cystic fibrosis. Curr. Pharm. Des. 18, 674–682 (2012).

12. Cutting, G. R. Modifier genes in Mendelian disorders: the example of cystic fibrosis: Modifiers of cystic fibrosis. Ann. N. Y. Acad. Sci. 1214, 57–69 (2010).

13. Williams, R. A., Mamotte, C. D. S. & Burnett, J. R. Phenylketonuria: an inborn error of phenylalanine metabolism. Clin. Biochem. Rev. 29, 31–41 (2008).

14. Dowell, R. D. et al. Genotype to Phenotype: A Complex Problem. Science 328, 469–469 (2010).

15. Kim, D.-U. et al. Analysis of a genome-wide set of gene deletions in the fission yeast Schizosaccharomyces pombe. Nat. Biotechnol. 28, 617–623 (2010).

16. Blomen, V. A. et al. Gene essentiality and synthetic lethality in haploid human cells. Science 350, 1092–1096 (2015).

17. Boutros, M. et al. Genome-wide RNAi analysis of growth and viability in Drosophila cells. Science 303, 832–835 (2004).

18. Hart, T. et al. High-Resolution CRISPR Screens Reveal Fitness Genes and Genotype-Specific Cancer Liabilities. Cell 163, 1515–1526 (2015).

19. Vu, V. et al. Natural Variation in Gene Expression Modulates the Severity of Mutant Phenotypes. Cell 162, 391–402 (2015).

20. Wang, T. et al. Identification and characterization of essential genes in the human genome. Science 350, 1096–1101 (2015).

21. Kamath, R. S. et al. Systematic functional analysis of the Caenorhabditis elegans genome using RNAi. Nature 421, 231–237 (2003).

22. Paaby, A. B. et al. Wild worm embryogenesis harbors ubiquitous polygenic modifier variation. eLife 4, (2015).

23. Edwards, M. D., Symbor-Nagrabska, A., Dollard, L., Gifford, D. K. & Fink, G. R. Interactions between chromosomal and nonchromosomal elements reveal missing heritability. Proc. Natl. Acad. Sci. 111, 7719–7722 (2014).

24. Hou, J., Tan, G., Fink, G. R., Andrews, B. J. & Boone, C. Complex modifier landscape underlying genetic background effects. Proc. Natl. Acad. Sci. 116, 5045–5054 (2019).

25. Galardini, M. et al. The impact of the genetic background on gene deletion phenotypes in Saccharomyces cerevisiae. Mol. Syst. Biol. 15, (2019).

26. Peter, J. et al. Genome evolution across 1,011 Saccharomyces cerevisiae isolates. Nature 556, 339–344 (2018).

27. Sadhu, M. J. et al. Highly parallel genome variant engineering with CRISPR-Cas9. Nat. Genet. 50, 510–514 (2018).

28. Michel, A. H. et al. Functional mapping of yeast genomes by saturated transposition. eLife 6, (2017).

29. Gangadharan, S., Mularoni, L., Fain-Thornton, J., Wheelan, S. J. & Craig, N. L. DNA transposon Hermes inserts into DNA in nucleosome-free regions in vivo. Proc. Natl. Acad. Sci. U. S. A. 107, 21966–21972 (2010).

30. van Opijnen, T. & Levin, H. L. Transposon Insertion Sequencing, a Global Measure of Gene Function. Annu. Rev. Genet. 54, 337–365 (2020).

31. Sharon, E. et al. Functional Genetic Variants Revealed by Massively Parallel Precise Genome Editing. Cell 175, 544-557.e16 (2018).

32. Hou, J. et al. The Hidden Complexity of Mendelian Traits across Natural Yeast Populations. Cell Rep. 16, 1106–1114 (2016).

33. Fournier, T. et al. Extensive impact of low-frequency variants on the phenotypic landscape at population-scale. eLife 8, e49258 (2019).

34. Harari, Y., Ram, Y., Rappoport, N., Hadany, L. & Kupiec, M. Spontaneous Changes in Ploidy Are Common in Yeast. Curr. Biol. 28, 825-835.e4 (2018).

35. Venkataram, S. et al. Development of a Comprehensive Genotype-to-Fitness Map of Adaptation-Driving Mutations in Yeast. Cell 166, 1585-1596.e22 (2016).

36. Johnson, M. S. et al. Phenotypic and molecular evolution across 10,000 generations in laboratory budding yeast populations. eLife 10, e63910 (2021).

37. Costanzo, M. et al. A global genetic interaction network maps a wiring diagram of cellular function. Science 353, (2016).

38. Costanzo, M. et al. Environmental robustness of the global yeast genetic interaction network. Science 372, eabf8424 (2021).

39. Martin, M. Cutadapt removes adapter sequences from high-throughput sequencing reads. EMBnet.journal 17, 10 (2011).

40. Li, H. & Durbin, R. Fast and accurate short read alignment with Burrows-Wheeler transform. Bioinforma. Oxf. Engl. 25, 1754–1760 (2009).

41. Giaever, G. et al. Functional profiling of the Saccharomyces cerevisiae genome. Nature 418, 387–391 (2002).

